# Visual performance fields in Saccadic Suppression of Image Displacement

**DOI:** 10.1101/2025.07.14.664707

**Authors:** Rosanne Timmerman, Antimo Buonocore, Alessio Fracasso

**Affiliations:** Psychology, School of Humanities, University of Dundee, Social Sciences and Law, Scotland; School of Psychology and Neuroscience, University of Glasgow, Glasgow, Scotland; Department of Educational, Psychological and Communication Sciences, Suor Orsola Benincasa University, Naples, Italy, 80135; Department of General Psychology, University of Padua, Padua, Italy, 35131

**Author notes:** Corresponding author, Alessio Fracasso Via Venezia 13 35121, University of Padua, Padua, Italy. shared senior authorship (Antimo Buonocore, Alessio Fracasso).

**Keywords:** saccadic suppression of image displacement, SSID, performance field, anisotropies, eye movements, saccade

## Abstract

Visual perception is not homogeneous throughout the visual field. Performance is generally better along the horizontal meridian compared to the vertical meridian, and in the lower compared to the upper visual field. These asymmetries in visual performance are reflected in structural asymmetries in early visual cortex.

When exploring a visual scene, eye movements occur continuously, with visual perception resulting from a tight interplay between the visual as well as the oculomotor systems. Literature on visual performance across visual fields during saccades is limited, but existing studies show that perceptual performance during saccades is indistinguishable between the upper and the lower visual fields, or altogether better in the upper visual field compared to lower.

In the current exploratory study, we asked participants to detect the direction of target displacement across visual fields, while performing a saccade as well as at fixation.

During fixation and saccade viewing conditions, performance on the task was better along the horizontal compared to the vertical meridian. However, we did not observe a robust difference in performance between the lower and upper visual field, neither at fixation nor when participants were requested to perform saccades. We interpret our results based on known behavioural and neural anisotropies, as well as considering evolutionary approaches to the perception-action cycle.

## Introduction

Visual perception is not homogeneous across the visual field (Carrasco et al., 2001; Fuller and Carrasco, 2009; Zito et al., 2016a; Barbot et al., 2021). Depending on where a visual stimulus appears, visual performance can differ substantially. For example, while fixating, performance on orientation discrimination tasks is generally higher for stimuli presented along the horizontal meridian compared to the vertical meridian (Horizontal Vertical Anisotropy, HVA). Additionally, performance is better in the lower visual field compared to the upper visual field (Vertical Meridian Asymmetry, VMA; (Carrasco et al., 2001; Fuller and Carrasco, 2009; Zito et al., 2016; Barbot et al., 2021).

These perceptual anisotropies are coupled with known cortical magnification anisotropies in primary visual cortex, suggesting how a larger portion of surface area in primary visual cortex is dedicated to processing the horizontal versus the vertical meridian and the lower compared to the upper visual field (Himmelberg et al., 2022; Kupers et al., 2022; Himmelberg et al., 2023).

Visual field anisotropies (or asymmetries) have been probed behaviourally at fixation using perceptual discrimination tasks such as orientation discrimination, target detection and contrast discrimination (Carrasco et al., 2001; Kristjansson and Sigurdardottir, 2008; Barbot et al., 2021). Moreover, they have been reported also in the context of deployment of attentional resources (He et al., 1996), and visual illusion (Rubin et al., 1996).

However, in daily life, humans rarely stare at one point for long, but make frequent eye movements to scan their surroundings. It is not clear whether visual field asymmetries normally observed at fixation persist also when saccades are involved, and whether the known pattern of visual anisotropies (HVA and VMA) would emerge also in this context.

Literature on visual field anisotropies while participants are requested to perform saccades is limited. Fracasso and colleagues (2023) showed that close to saccade onset, participants perform better in an orientation discrimination task in the upper visual field compared to the lower visual field. Furthermore, Liu and colleagues (Liu et al., 2024), showed a significant reduction in VMA reduction when participants were asked to perform saccades compared to fixation. As a third example, Grujic et al., (2018) measured the errors in perisaccadic localization of briefly presented targets when participants were requested to perform horizontal or vertical saccades. The authors observed significant differences in performance between upward and downward saccades (Grujic et al., 2018).

Although the tasks adopted by these studies are different, spanning between orientation discrimination and target localization, it is interesting to note how these studies are suggestive of an overall VMA reduction or inversion during saccades compared to fixation (Grujic et al., 2018; Hafed, 2018; Fracasso et al., 2023; Liu et al., 2024). Grujic and Colleagues (2018) suggest how effects in perceptual localization along the vertical meridian might depend on neural circuits that asymmetrically represent the upper and lower visual fields, as for example in the superior colliculus (SC) in the midbrain (Grujic et al., 2018; Hafed & Chen, 2016). The superficial and deeper layers of the SC are organized retinotopically with corresponding visual and oculomotor maps, and it houses motor neurons controlling eye muscles and generating saccades in humans (Hafed et al., 2023) and non-human primates alike (Wurtz and Goldberg, 1972; Mohler and Wurtz, 1976; Wurtz and Albano, 1980). Interestingly, visual and motor neuronal responses in SC shows opposite anisotropies compared to primary visual cortex along the vertical meridian, showing faster and stronger neuronal responses for stimuli presented in the upper visual field compared to the lower visual field (Hafed and Chen, 2016; Hafed 2018; Fracasso et al., 2023; Hafed et al., 2023).

In the current study we measure visual field anisotropies at fixation and while participants are asked to perform saccades. We implemented the saccadic suppression of image displacement paradigm (SSID), a task in which participants are required to discriminate the direction of a visual target that is displaced while the eyes are moving toward it. In SSID, a participant’s ability to discriminate the direction of target displacement is severely impaired when the eyes are moving compared to fixation (Bridgeman et al., 1975; Deubel et al., 1996; Deubel et al., 1998; Deubel et al., 2002). In the original study by Bridgeman et al. (1975), the authors were interested in studying the influence of an anticipatory signal generated by the oculomotor system to the visual system (corollary discharge), to account for the upcoming retinal shift introduced by the following eye movement.

In this exploratory study, we aim to assess whether and how anisotropies observed at fixation (HMA and VMA) translate in the context of saccades. Specifically, given the existing results reported in the literature, we expect an overall reduction in VMA (smaller or no difference between the upper and lower VPF) and a largely unaltered HMA (better performance along the horizontal compared to the vertical meridian).

## Methods

### Participants

A total of 25 participants between the age of 18 and 39 took part in the measurements (19 females). One participant was excluded because they showed non-collaborative behaviour. All participants reported normal or corrected-to-normal vision. The experiment was approved by the local ethics committee at the College of Medical, Veterinary and Life Sciences. Written informed consent was obtained, in accordance with the 1964 Declaration of Helsinki. Participants received compensation of £6 per testing hour. Each participant took part in a variable number of experimental sessions (2 or 3). A variable number of blocks (between 10 and 29 blocks) were completed by each participant. The variability in the number of blocks performed is due to scheduling restraints, as some participants did not return for the third session. Experimental sessions took place on non-consecutive days, each taking about 60 minutes.

### Experimental design

We measured the performance on a target displacement task in the periphery using a two-by-two within-participants design to study the influence of *meridian* (horizontal or vertical target movements) and *viewing condition* (fixation or saccades). For each experimental condition, four displacements were adopted: -1.7, -0.5, 0.5 and 1.7 degrees of visual angle (dva). Positive values indicate displacements away from the screen centre. Negative values indicate displacements towards the screen centre.

### Apparatus

Participants were placed in a chin- and forehead rest to ensure the stability of the head. Responses were given by pressing keys on a standard keyboard. Stimuli were presented on a 24-inch LCD monitor (1920 x 1024 pixels) with a refresh rate of 144 Hz. Display luminance was linearized, and gamma corrected. A screen cover was placed over the screen to create a square aperture (see Figure 1). We opted for this experimental setup as visual references affect how we (mis)localize objects along the perisaccadic interval. More specifically, visual references directly affect perceived location and visual compression (Lappe et al., 2000). Furthermore, visual references can modulate performance in SSID (Deubel et al., 1998).

**Figure 1.**
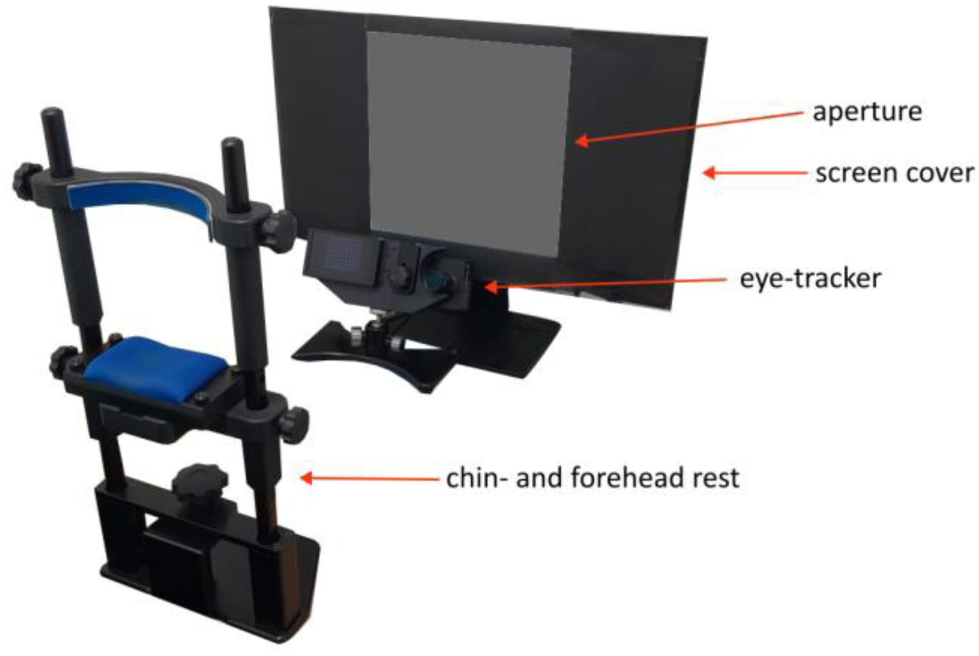
Experimental set-up. Participants sat in a dimly lit room and took position in the chin- and forehead rest, with the eye-tracker camera focused on their left eye. The screen was partly covered with black cardboard to create a square aperture.

Eyes were horizontally and vertically aligned with the centre of the screen at a distance of 64.5 cm. Eye position was sampled using an Eyelink 1000 (SR Research, Ltd., Ottawa, ON), acquiring data at 1000 Hz. A five-point calibration in a square-shaped pattern was performed at the beginning of each experimental block. Both eyes were tracked for the duration of the experiment. The experiment was programmed in Matlab (R2021a, The Math Works, Inc., 85 Natick, MA), with the Psychtoolbox (Brainard and Vision, 1997) and the Eyelink toolbox (Cornelissen et al., 2002).

### Experimental procedure

The experiment consisted of three parts. During the first part, we measured an estimate of individual saccade latency separately for vertical and horizontal saccades. This individual reaction time was used later in the main part of the experiment to ensure an equivalent delay in displacement onset between the saccade and fixation condition. During the saccadic latency test, a starting fixation point of .35 dva was presented in the middle of the screen. Participants sat in a dimly lit room and were instructed to fixate on the starting fixation point and to press the spacebar to start each trial. After a variable delay (1 - 1.5s) the starting fixation point disappeared from the centre of the screen and reappeared at 10 dva in the periphery – the target fixation point. For horizontal saccades, the target fixation point could be presented randomly on the left or right, along the horizontal meridian. For vertical saccades, the target fixation point could be presented randomly on the upper or lower part of the screen, along the vertical meridian. Participants were instructed to perform an eye movement towards the target fixation point as accurately and as rapidly as possible. After each trial, we derived an estimate of saccade reaction time. Saccade latency towards the left/right (horizontal meridian) and upper/lower (vertical meridian) was tested in two separate blocks of 40 trials each. At the end of each training block, we derived the median saccade reaction time for horizontal and vertical saccades, separately. The saccade latency test lasted approximately 10 minutes.

Following the saccade latency test, the participant performed a training session consisting of 15 trials to become familiar with the experimental procedure. Participants were instructed to fixate on the starting fixation point and to press the spacebar to start each trial. After a variable delay (1 -1.5 s) the starting fixation point disappeared from the centre of the screen and reappeared at 10 dva in the periphery (left or right, along the horizontal meridian - the first movement). After a variable interval, between 80% and 120% of the individual participant’s median saccade reaction time in the horizontal condition, the target was displaced with -1.7, -0.5, 0.5 and 1.7 dva relative to the target fixation point, along the horizontal meridian. Throughout the manuscript, we use the term ‘first movement’ when referring to the shift of the initial fixation dot towards the periphery, and the term ‘displacement’ when referring to the second shift of the peripheral fixation dot.

During the third part participants performed the main task. Before each block, participants were informed of the task condition they would be taking part in (horizontal or vertical meridian) and whether to keep fixation (fixation condition) or follow the dot with their eyes (saccade condition). Each trial started with a black fixation point with a size of .35 dva appearing in the centre of the screen. Participants were instructed to fixate and press the spacebar to start each trial. After a variable interval (750 - 1250 ms) the fixation point moved either 10 dva to the left or to the right (horizontal condition), or up or down (vertical condition) – the first movement. During ‘fixation’ blocks, participants were instructed to keep stable fixation in the centre of the screen. During ‘saccade’ blocks, participants were instructed to move their eyes as accurately and rapidly as possible towards the displaced target. Shortly after the first movement of the target, the target moved again, 0.5 or 1.7 dva along or against the direction of the first target movement (displacement, see Figure 2 for a graphic representation of the trial procedure in the saccade and fixation conditions).

**Figure 2.**
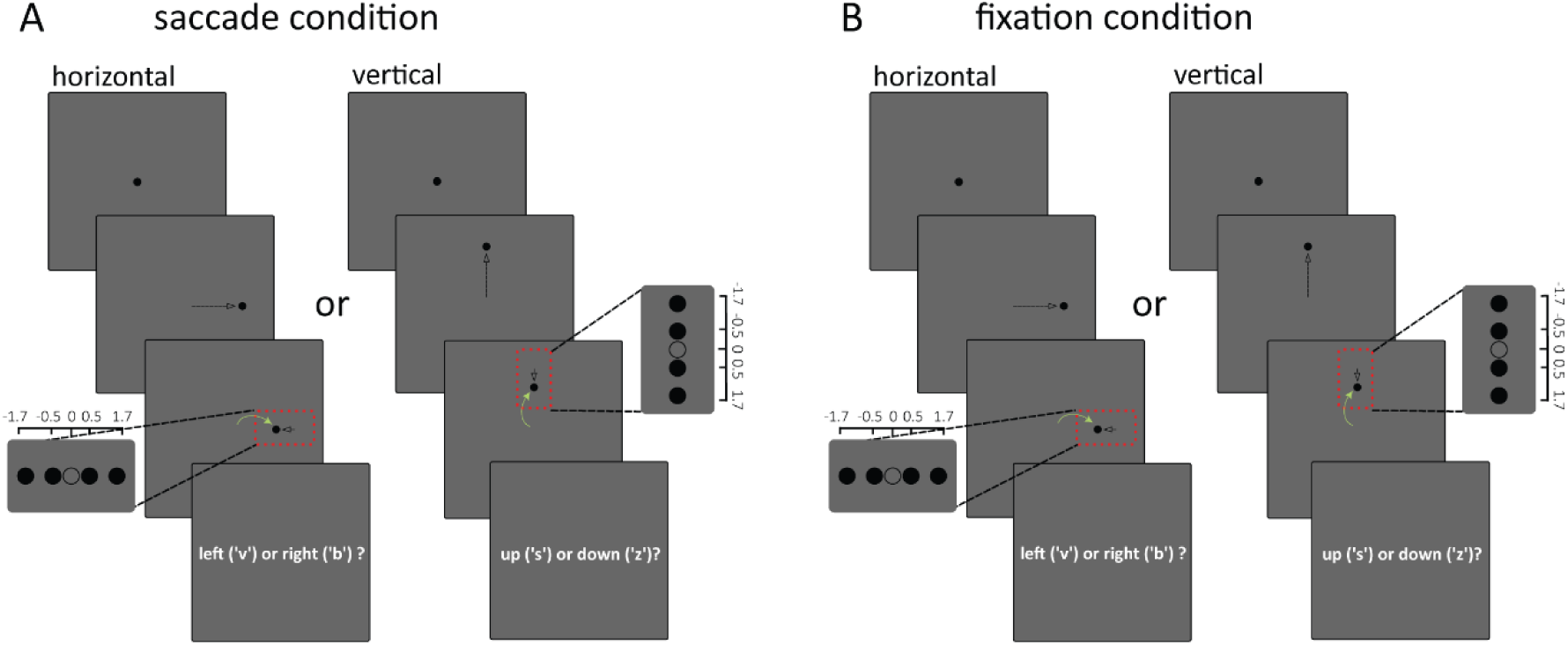
Experimental design **(A)** Each trial started with a black fixation point with a size of .35 dva appearing in the centre of the screen. Participants were instructed to fixate and press the spacebar to start the trial. After a variable interval (between 750 and 1250 ms) the fixation point moved either 10 dva to the left (or right, horizontal condition), or up (or down vertical condition) – the first movement. Participants were instructed to move their eyes as accurately and rapidly as possible towards the target fixation point (green arrow). Shortly after the first movement of the target, the target moved again, 0.5 or 1.7 dva along or against the direction of the first movement. The onset of the displacement was contingent on the experimental condition. In the saccade condition, the displacement was presented in the first monitor flip immediately after saccade detection; therefore, the displacements only took place while the eyes were ‘in flight’. After each trial, participants were instructed to press a key on the keyboard corresponding to the direction of the displacement of the target. The direction of the first movement and the displacement (up;down / left;right) was randomized for each trial. **(B)** The trial presentation during the fixation condition was identical to the saccade condition, the only differences being that participants were instructed to keep stable fixation in the centre of the screen throughout the trial, and the displacement was randomly presented between 80% and 120% of the individual participant’s median saccade reaction time.

The onset of the displacement was contingent on the experimental condition. On ‘saccade’ blocks, saccades were detected by means of a gaze-contingent algorithm (Schweitzer and Rolfs, 2020). The displacement was presented in the first monitor flip immediately after saccade detection. This ensured that the displacements only took place while the eyes were ‘in flight’. On ‘fixation’ blocks, the displacement was presented between 80% and 120% of the individual participant’s median saccade reaction time in the corresponding condition (horizontal or vertical). For example, if someone’s median saccade reaction time was 170ms for horizontal saccades, the displacement would randomly occur between 136 and 204 milliseconds (Fabius et al., 2016; Fracasso and Melcher, 2016; Buonocore et al., 2017; Fabius et al., 2020; Taylor et al., 2024).

After each trial, participants were asked to press a key on the keyboard corresponding to the direction of the displacement of the target (‘v’ for left, ‘b’ for right for the ‘horizontal’ condition, ‘s’ for up and ‘z’ for down for the ‘vertical’ condition). A written reminder of the response mapping was presented onscreen after each trial. The direction of the first movement and the displacement was randomized for each trial (see Figure 2).

Each block consisted of 45 trials and lasted for approximately 3 minutes. Drift correction was applied every 23 trials.

During day 1, participants would complete the saccadic latency tests, the training, and several blocks of the main task. On days 2 and 3, participants continued with the main task, without a training session (Fracasso et al., 2010; Kaunitz et al., 2011; Melcher and Fracasso, 2012; Fracasso et al., 2013; Kaunitz et al., 2013; Kaunitz et al., 2014; Fabius et al., 2019; Taylor et al., 2024).

## Data Analysis

### Data processing and analysis: psychometric fitting

We used Matlab2021a (www.themathworks.com) to combine the eye tracking and behavioural data. Data wrangling, filtering, and analysis were carried out in R v4.2.0. Data were filtered based on several criteria: 1) whether participants were correctly performing eye movements or kept fixation in the corresponding experimental blocks, 2) for the ‘saccade’ condition, whether the first saccade was in the correct direction (left/right/up/down) 3) whether the displacement occurred while the eyes were ‘in flight’ (between saccade onset and saccade onset + saccade duration), 4) the duration of the first saccade being shorter than 95 ms and lastly, 5) whether the amplitude of the first saccade was at least 6.5 dva. One participant was excluded, and after applying the above filters we removed 18.9% of trials for the vertical condition at fixation and 22.3% of trials were removed in the saccade condition. Regarding the horizontal condition, 10.9% of trials were excluded from the fixation condition and 14.3% of trials were removed from the saccade condition. Most of the exclusions in the saccade condition were due to displacement onset being outside the range of saccade execution, either slightly before or after saccade onset. As for the fixation condition, we excluded trials if a saccade larger than 0.5dva was detected within -80ms to +80ms around the displacement onset.

We used a Generalized Linear Model (GLM) to analyse the individual participant data. We fit the proportion of responses along the direction of the first movement as a function of the displacement direction and size (-1.7, -0.5, 0.5, 1.7) using a logistic regression model for each participant, visual field (left, right, upper, or lower) and viewing condition (saccade and fixation).

We computed the probability of giving a response along the direction of the first movement, Pr(*y*_*i*_ = 1), as a function of the size and direction of the displacement (*x*).

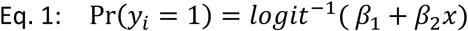

Specifically, we map the values to the range (0,1) using the following:

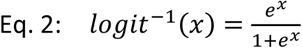

From the output of this model, the bias, slope, were extracted for each participant and for each of the above-mentioned conditions. The bias was calculated by dividing the first coefficient (the intercept) by the second coefficient on the Generalized Linear Model. This represents the point where the predicted probability is 0.5 (Point of Subjective Equality; PSE). In other words, the point when participants were unable to discriminate between the direction of the displacement (along the direction of the first movement or in the opposite direction). A negative bias indicated a bias *along* the direction of the first movement, whereas a positive bias indicates a bias *against* the direction of the first movement (Fig. 3a).

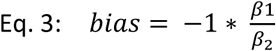

**Figure 3.**
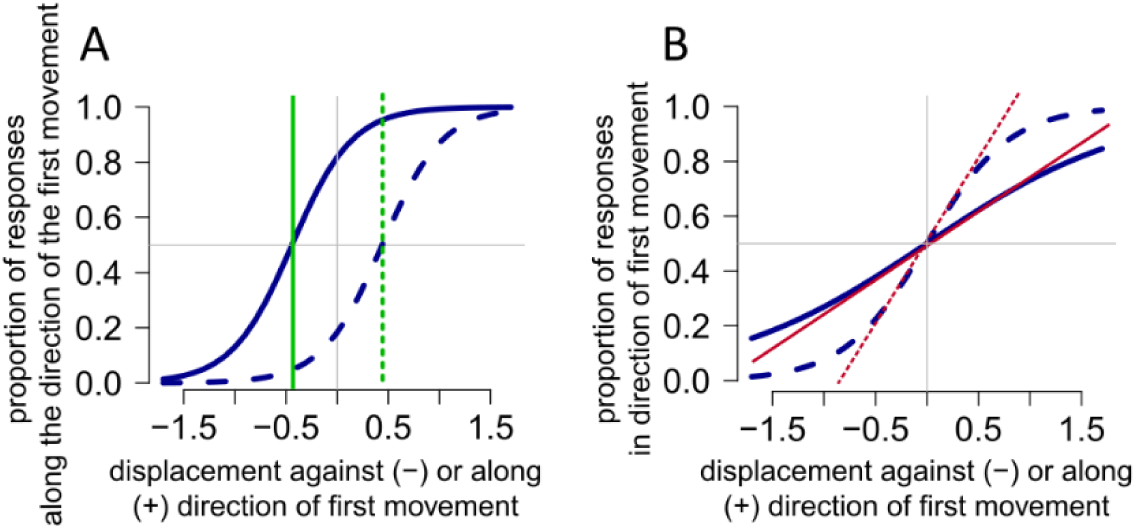
Psychometric curve fitting **(A)** Differences in bias. The continuous blue line shows a response pattern with a relatively steep slope and a systematic bias in reporting the direction of the displacement along the direction of the first movement (continuous green line, negative bias), whereas the dashed blue line shows a systematic bias in reporting the direction of the displacement against the direction of the first movement (dashed green line, positive bias). See main text for further details. **(B)** Examples of differences in slopes. The continuous blue line depicts a response pattern with a shallow slope (small slope, closer to 0), indicating a relatively poor performance in discriminating the displacement direction. The dashed blue line shows a steeper slope (larger slope), indicating a relative better performance.

The slope of the logistic regression curve is steepest at this PSE and was calculated by dividing the second coefficient by four and by taking the absolute. *β*_2_/4 is the maximum difference in Pr(*y*_*i*_ = 1) corresponding to the rate of change per unit x.

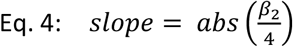

The slope refers to the rate of change of the proportion of responses along of the direction of the first movement. A steeper slope (larger slope) indicates better performance on discriminating the displacement, a shallower slope (smaller slope, closed to 0) indicates poorer performance (Fig. 3b). The reported equations and a detailed description of their interpretation can be found in *Data Analysis Using Regression and Multilevel/Hierarchical Models*, chapter 5, Logistic regression, section 5.2: Interpreting the logistic regression coefficients (Gelman and Hill, 2007).

For each participant and viewing condition we computed the average slope and bias for the horizontal (average between left and right visual fields) and vertical meridian (average between upper and lower visual fields).

### Statistical analysis

We used paired t-tests to compare the differences in slopes and bias between the following conditions: 1) horizontal versus vertical meridian for saccade condition, 2) horizontal versus vertical meridian for fixation condition, 3) upper versus lower visual field for saccade condition, and lastly, 4) upper versus lower visual field for fixation condition. Paired t-tests were corrected using Bonferroni correction, corrected p values are reported. We report effect sizes for the significant comparisons (Cohen’s d).

We opted for comparing results within each condition (saccade and fixation) separately. We refrained from comparing different modalities (saccade VS fixation), as there are known large differences between these conditions in bias as well as slope. This is evident in the literature (Bridgeman et al., 1975; Deubel et al., 1996; Deubel et al., 1998; Deubel et al., 2002). Thus, we considered it redundant to report the results here as well. Moreover, especially for estimates of bias, the relative difference between modalities is not informative per se. The bias departure from 0 (either towards the positive or negative) is indicative of the direction of the biased response, either along or opposite with respect to the eye movement, or towards or away from fixation, thus providing a clear interpretation of the participants response tendency in the different conditions.

Lastly, we fitted Bayesian linear models and obtained Bayes factors from the BayesFactor package for R (Morey et al., 2015) comparing slope and bias estimates between experimental conditions. When describing our results, we use terminology discussed in (Lee and Wagenmakers, 2014) regarding evidence in favour or against the null hypothesis. More specifically, we use the terms ‘weakly in favour/against’, ‘moderately in favour/against’ and ‘strongly in favour/against’. These corresponding to Bayes factors of <3, 3–10 and > 10, respectively (Brunner et al., 2025).

### Clustering and individual differences

We performed a cluster analysis to capture variability in participants’ response patterns. We chose this purely data driven approach as it does not impose any constrains on the expected response patterns.

We adopted the hierarchical clustering implementation in R, using the function *hclust* and the maximum distance as a dissimilarity matrix. To determine the optimal number of clusters, we inspected the silhouette plots of the clustering solution (function *fviz_nbclust,* library *factoextra*).

For any given entry in a set, the silhouette score measures how similar an entry is to its own cluster compared to other clusters. The silhouette score ranges from -1 to 1. Values above 0 indicate that the entry is well matched to its own cluster and poorly matched to neighbouring clusters. The opposite is true for values below 0. The silhouette score for the entire dataset is the average of the silhouette scores of all individual entries. The silhouette plot displays the measure of how close each point in one cluster is to points in the neighbouring clusters, providing a way to assess the optimal number of clusters (Dalby et al., 2024). An average silhouette plot was created for each of the four conditions (*meridian X viewing condition*) for a number of clusters ranging between 2 and 10 (Fig. 4).

**Figure 4.**
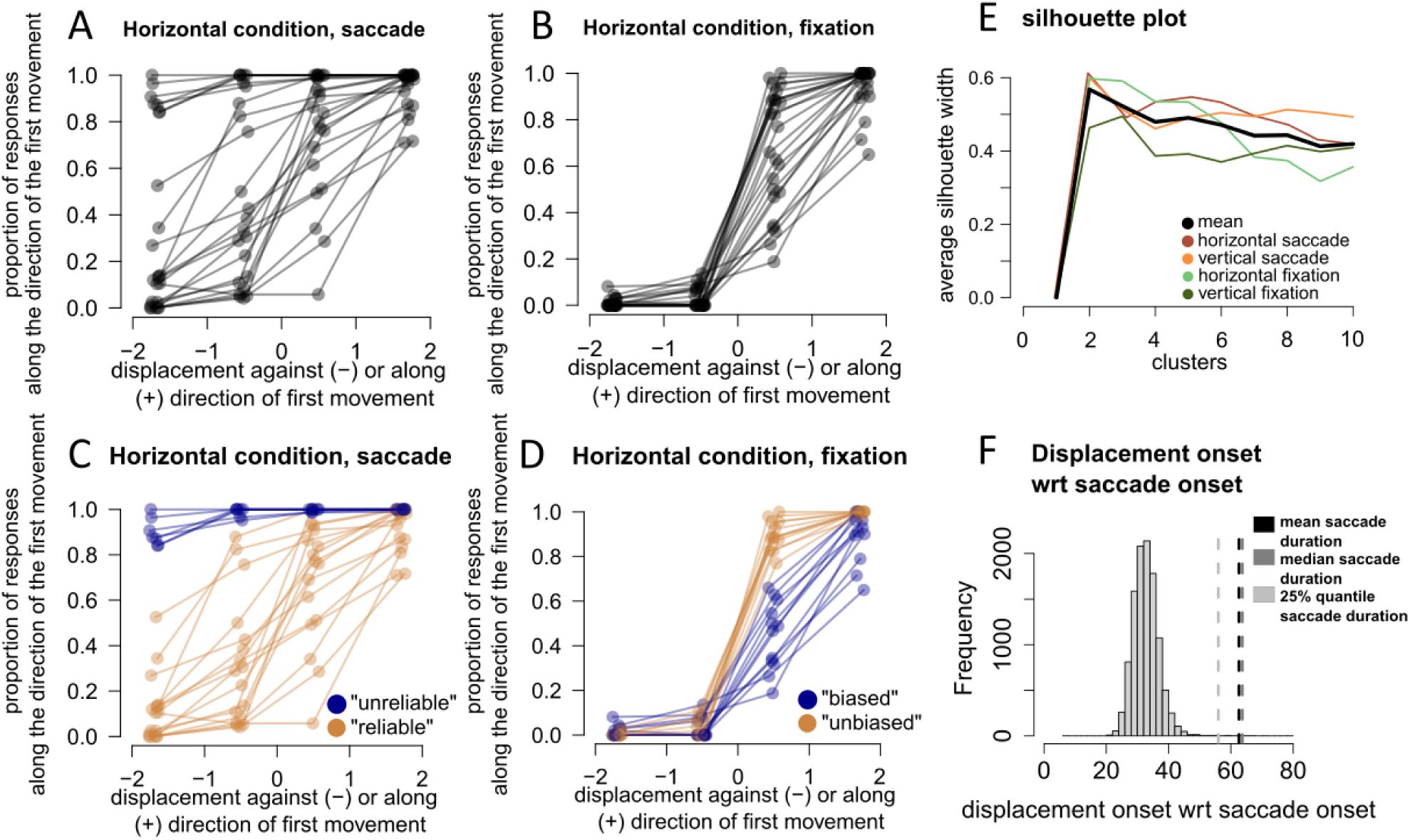
Inter-participant variability in responses during saccades and fixation **(A-D)** Each dot is the average response for the specific displacement of each participant (in the horizontal condition). The connecting line shows the response pattern per participant. **(A, B)** Responses of each participant during the saccade condition and fixation condition before clustering, respectively. **(C, D)** Same plots as in A, B but participants are divided into two different clusters, ‘unreliable’ and ‘reliable’. **(C)** ‘Unreliable’ participants have a strong bias to judge the displacement along the direction of the first movement, regardless of the direction of the displacement. **(D)** Despite the smaller inter-participant variability, when we applied the clustering algorithm to fixation condition, we observed 2 subgroups: ‘biased’ participants showed a tendency to judge the displacement against the direction of the first movement, thus closer to the fixation target than it really was. Although present, the bias in the fixation condition was considerably smaller than the unreliable and biased responses observed in the saccade condition. Importantly, participants that gave a systematically unreliable responses in the saccade condition were not the same that gave a biased response in the fixation condition (see section ‘Unreliable judgements in the saccade condition do not translate in the fixation condition’ in the main text) **(E)** Silhouette plot showing the silhouette width for each of the four conditions (meridian X viewing condition) for different clusters. The mean of the four conditions shows that the width is highest at two clusters. **(F)** The histogram shows the distribution of displacement onset of all trials included in the analysis (saccade condition). Dashed lines indicate the mean, median and the 25% quantile saccade reaction time. Displacement onsets are well within the range of saccade duration, ensuring that the displacements only took place while the eyes were ‘in flight’.

In Figure 4, panels A-D provide a description of the steps adopted in our clustering approach, as well as the outcome, using data from horizontal saccades as an example.

## Results

We aimed at testing behavioural performance along different visual fields in a displacement discrimination while participants were requested to perform eye movements or at fixation.

### Clustering and inter-participant variability: unreliable responses in the saccade condition

Participants responses in the saccade condition (Fig. 4a) showed considerably more variability than in the fixation condition (Fig. 4b). We inspected the average silhouette of the four conditions (black line in fig. 4e) and determined 2 as the optimal number of clusters (average width of .57). While inspecting the clustering outcome, we observed a clear distinction in the reliability of estimates between two groups of participants during the saccade condition. An unreliable group would consistently judge the displacement along the direction of the first movement, and therefore along direction of eye movement (Fig. 4c). As in this group the bias and slope estimates were attainable only by extrapolating the fits, making any precise estimation unreliable, we used the same terminology to refer to this group of participants (‘unreliable’ vs ‘reliable’).

When we applied the clustering algorithm to the fixation condition, we observed 2 subgroups: ‘biased’ participants showed a tendency to judge the displacement against the direction of the first movement, thus closer to the fixation target than it really was (Fig. 4d). Although present, the bias in the fixation condition was considerably smaller than the one observed in the unreliable group, in the saccade condition.

### Unreliable judgements in the saccade condition do not transfer to the fixation condition

Seven participants were part of the ‘unreliable’ group in the saccade condition. This sub-group of participants gave consistently biased responses along all saccade conditions (horizontal and vertical), with an average overlap of 83% across conditions, that is: a participant assigned to the ‘unreliable’ group in one experimental condition (e.g. saccades towards the left visual field) was also assigned to the same group (‘unreliable’) in a different experimental condition (e.g. saccades towards the right visual field).

For the seven participants that were part of the ‘unreliable’ group in the saccade condition, bias and slope estimates were attainable only by extrapolating the fits, thus making any precise estimation unreliable. For this reason, we did not analyse the data further from these seven participants in the saccade condition, limiting the number of participants to 17.

In Figure 5 we show the split between unreliable and reliable participants from the saccade condition (Fig. 5a). Furthermore, we show the same split (based on saccade condition) but applied to the fixation condition (Fig. 5b). As shown, participants who gave systematically unreliable responses in the saccade condition were not similarly unreliable in the fixation condition.

**Figure 5.**
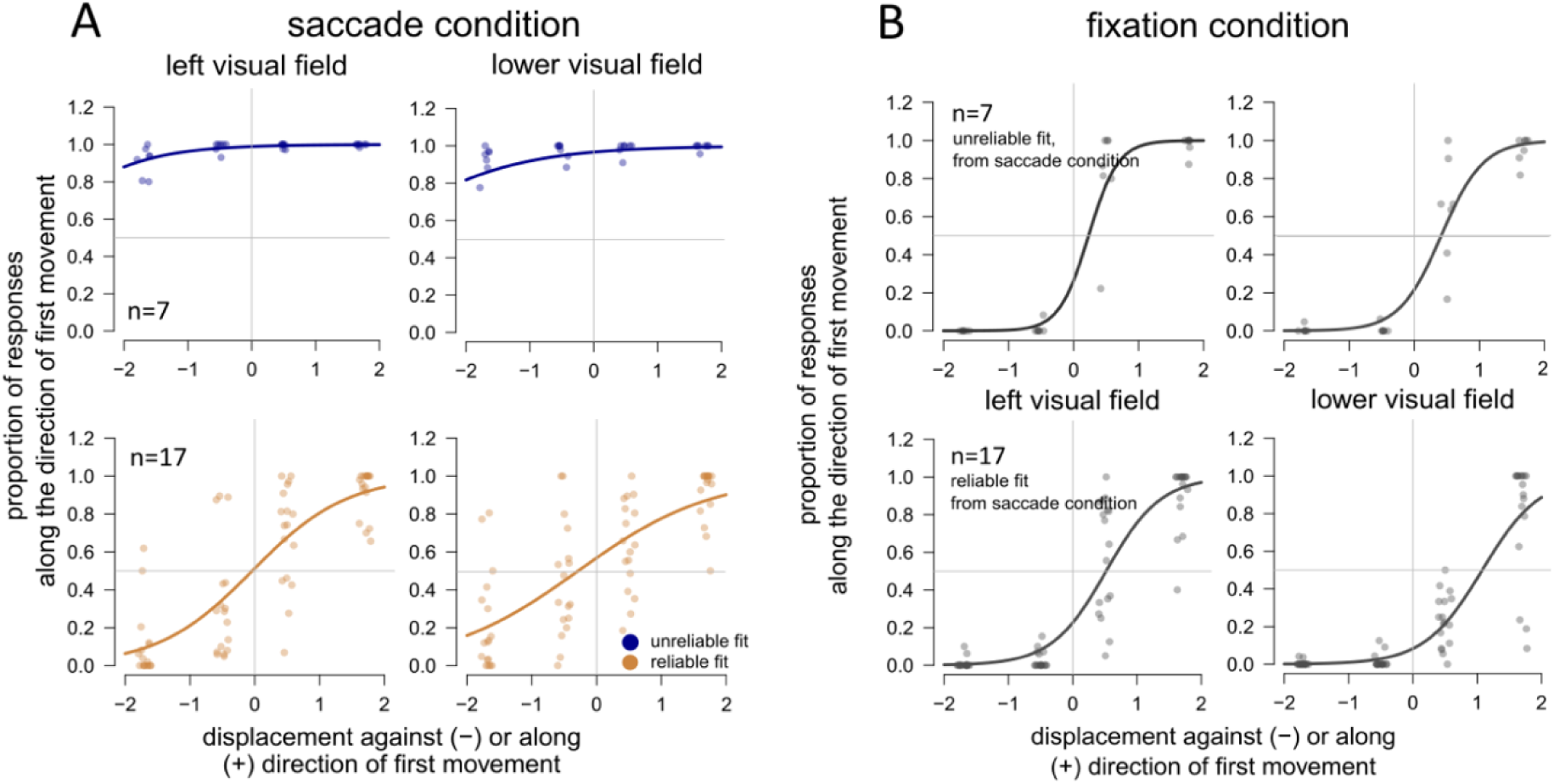
Clustering of participants per viewing condition. We show the average fit per cluster in the left and lower visual field, resulting from the GLM. **(A, B)** The responses of the clusters are similar across different visual fields. **(A)** There is a clear distinction between the ‘unreliable’ and ‘reliable’ cluster in the saccade condition, in which the ‘unreliable’ group would consistently judge the displacement along the direction of the first movement, and therefore along direction of eye movement. **(B)** When we apply the clustering obtained from the saccade condition to the fixation condition, responses are more uniform, indicating that unreliable judgements in the saccade condition do not transfer to the fixation condition.

Saccade properties do not differ between reliable and unreliable clusters. Amplitude between unreliable (M=8.35) and reliable (M=8.16) cluster did not differ statistically, t(11.7) = 1.78, p = 0.101, neither did saccade duration (M=67.13, unreliable; M=69.19, reliable), t(10.6) = -0.95, p = 0.365. There also was no difference in peak velocity (M=400dva/s unreliable, M= 406 dva/s reliable), t(12.5) = -0.32, p = 0.753. Lastly, there was no difference (M= 162 reliable, M=152 unreliable) in saccadic reaction time: t(11.9) = 1.46, p = 0.169.

Extrapolation was not an issue for data acquired at fixation. Moreover, responses at fixation were more homogeneous between participants, and we observed a general bias to report displacements against the direction of the first movement (see Fig. 5b). This represents an ‘inward’ bias where displacements are judged as being closer to the fixation point than physically presented. Previous literature shows a similar inward bias, although it is important to note that most of these studies looked at perceived location of a target, not displacement (Musseler et al., 1999; Eggert et al., 2001; Sheth and Shimojo, 2001; Awater and Lappe, 2006; Brenner et al., 2006; Brenner et al., 2008).

Nonetheless, as a further control, we applied the same clustering analysis to the fixation condition and observed comparably less overlap between conditions at fixation (horizontal and vertical, with an average between-participants overlap of 63% across conditions: a participant assigned to the ‘biased’ group in one experimental condition was not assigned to the same group in a different experimental condition). Considering: i) the lower overlap between experimental conditions at fixation and that ii) extrapolation was not an issue at fixation, we did not split the participants at fixation in ‘biased’ and ‘unbiased’ for the following analyses and used the complete dataset of 24 participants for the results at fixation. For visualization purposes only, we show the results split based on clustering outcome at fixation in Fig. 4d.

To summarize, following the clustering analysis we removed 7 participants in the saccade condition, which left us with 17 participants in the saccade condition and 24 participants in the fixation condition.

### Fixation condition, comparing bias and slopes for HMA and VMA

Individual fits resulting from the GLM were calculated for each participant and condition and are shown in Fig. 6 and 7. In the fixation condition participants were significantly more precise in detecting displacements in the horizontal plane in comparison to the vertical plane, *t*(23) = 4.62, corrected *p*<0.001, *d* = .94 (uncorrected *p* < 0.001; Fig. 6a,b left panel). No difference in performance was found between the lower and upper visual field, *t*(23) = -0.753, *ns*, corrected *p* = 1.00 (uncorrected *p* =0.470; Fig. 6c,d left panel). We found a significant difference in bias between movement in the horizontal and vertical plane (Fig. 6b, right panel). Although there was an inward bias present in both the horizontal and vertical condition, displacements along the vertical plane were judged closer to the fixation point compared to the horizontal plane, *t*(23) = -4.29, *p*<0.001 (uncorrected *p* < 0.001). No difference in bias was found between the upper and lower visual field, *t*(23) = 1.06, corrected *p* = 1.00 (uncorrected *p* =0.300, Fig. 6d, right panel).

**Figure 6.**
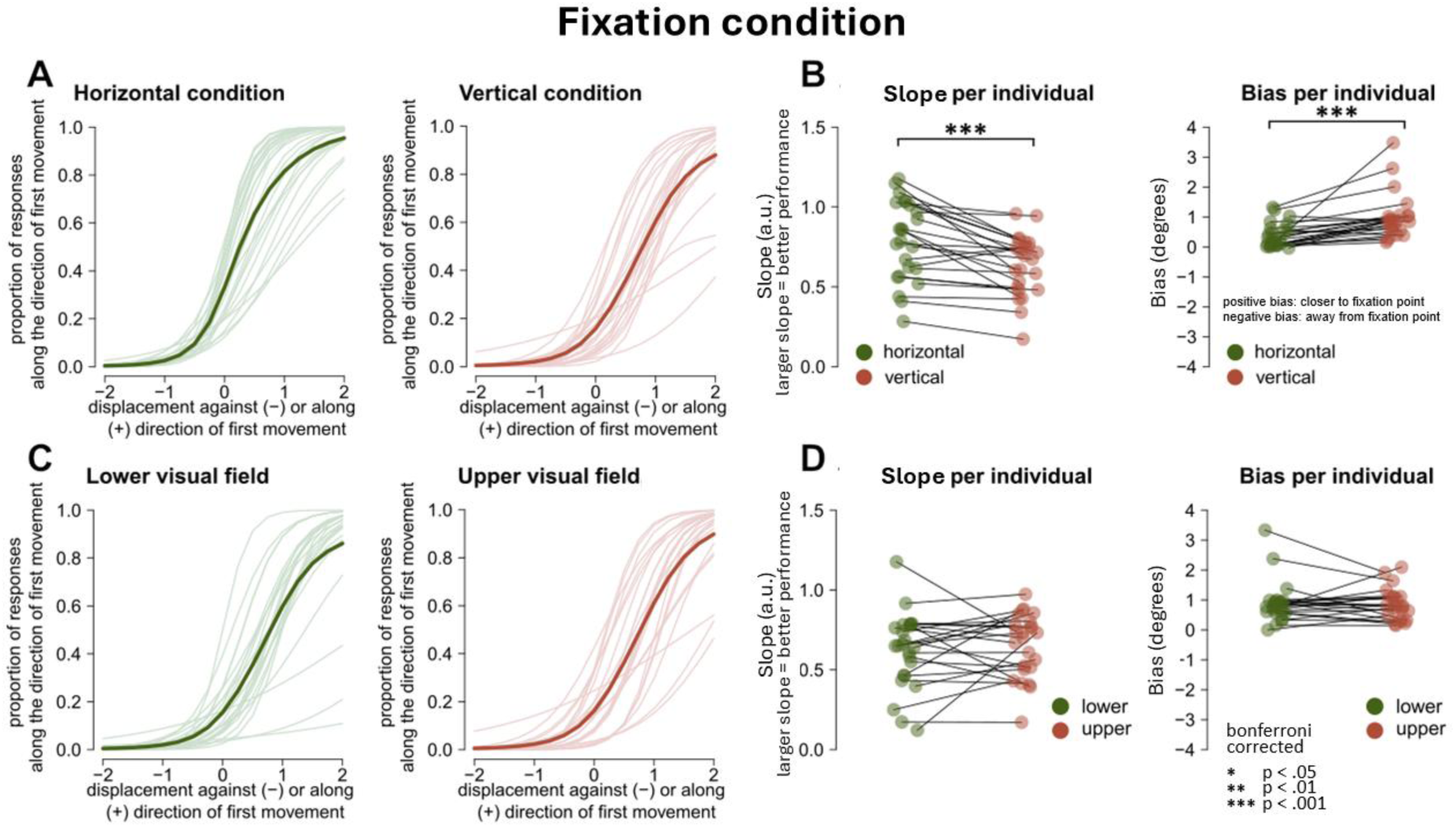
Individual fits, slopes and biases during fixation. **(A)** Individual fits from GLM in the horizontal and vertical condition. Thick line: average fit across participants **(B)** Slope (left) and bias (right) per participant in horizontal and vertical condition. **(C)** Individual fits from GLM in the lower and upper visual field. **(D)** Slope (left) and bias (right) per participant in upper versus lower visual field.

**Figure 7.**
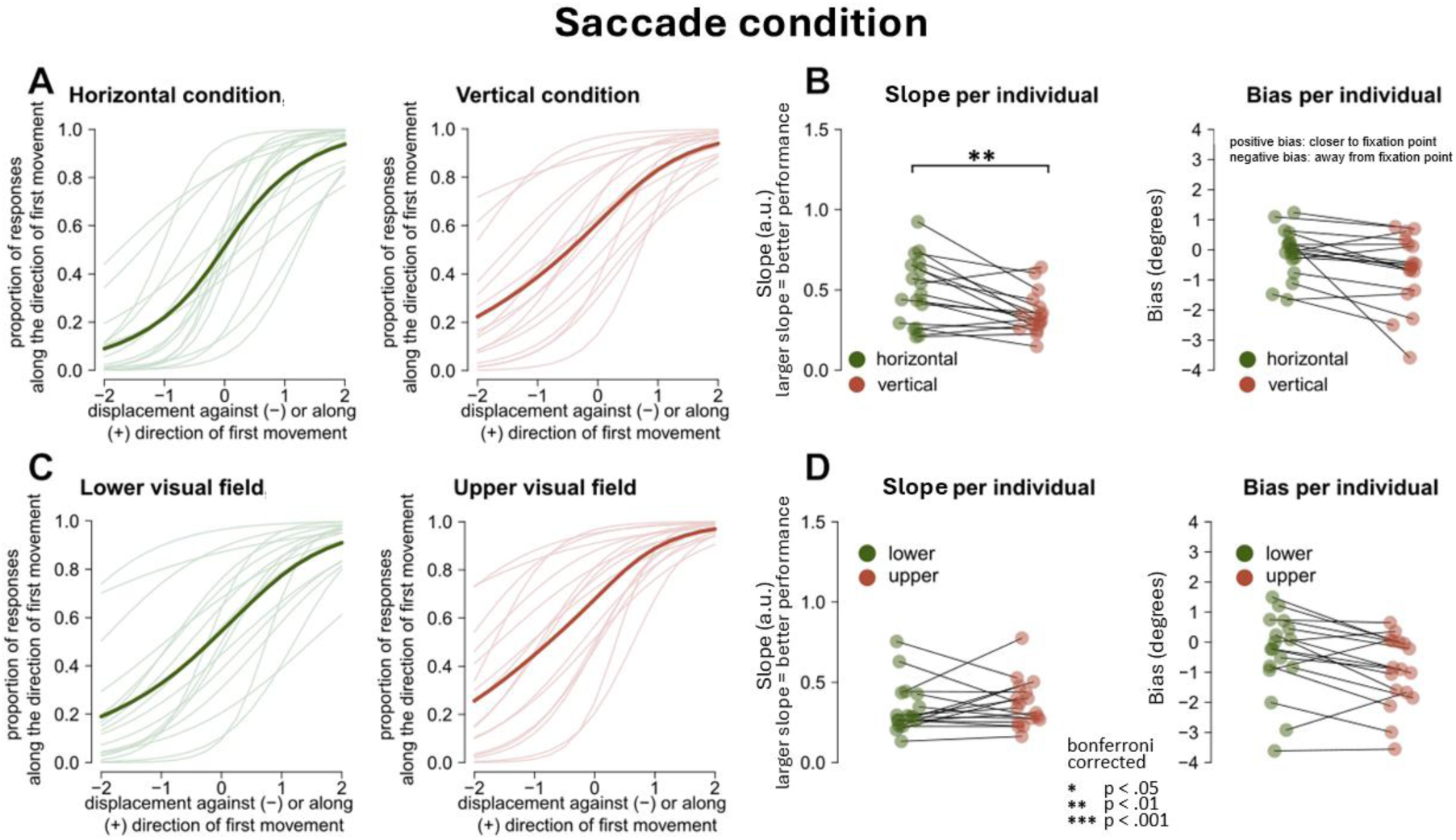
Individual fits, slopes and biases during saccade condition. **(A)** Individual fits from GLM in the horizontal and vertical condition. Thick line: average fit across participants **(B)** Slope (left) and bias (right) per participant in horizontal and vertical condition. **(C)** Individual fits from GLM in the lower and upper visual field. **(D)** Slope (left) and bias (right) per participant in upper versus lower visual field.

### Saccade condition, comparing bias and slopes for HMA and VMA

In the saccade condition, the slopes in the horizontal condition are steeper than the slopes in the vertical condition (Fig. 7a), indicating better performance in the former condition. Indeed, a paired t-test showed that participants were more precise at detecting displacements in the horizontal plane in comparison to the vertical plane, *t*(16) = 3.53, corrected *p* = 0.010, *d* = .86 (uncorrected *p* = 0.003) (Fig. 7b, left panel). The average fit of the lower and upper visual field looks very similar (Fig. 7c), and the paired t-test confirmed that there was no significant difference in slope between the lower and upper visual field, *t*(16) = -0.70, corrected *p* = 0.490*, ns,* (uncorrected *p* = 0.300 ; Fig. 7d, left panel).

In the saccade condition, there appears to be a trend in which the bias was stronger for the vertical condition in comparison to the horizontal plane, *t*(16) = 2.52, *ns, p* = 0.090*, d* = .61 (uncorrected *p* = 0.020), and in the upper visual field in comparison to the lower visual field, *t*(16) = 2.75, *ns, p* = 0.056*, d* = .67 (uncorrected *p* = 0.014); Fig 7b-d), but the tests did not survive Bonferroni correction.

Although this trend did not survive correction for multiple comparisons, it is interesting to note the following: if we consider a bias of 0 as veridical performance, then having a departure from 0 would indicate a less veridical percept. A stronger bias for the vertical condition compared to the horizontal condition, and stronger bias for upper versus lower visual field would indicate a more veridical percept along the horizontal compared to the vertical condition, and a more veridical percept for the lower compared to the upper visual fields. This pattern is compatible with the HVA and VMA observed at fixation in perceptual discrimination (Carrasco et al., 2001; Fuller and Carrasco, 2009; Zito et al., 2016; Barbot et al., 2021), and goes in the opposite direction compared to the perisaccadic modulation shown in Fracasso et al. (2023).

### Bias and slope estimates along visual fields and viewing conditions

In the following, we report the individual estimates of slope (Fig. 8a-c) and bias (Fig. 8d-e) across visual fields and viewing condition. In previous literature looking at perceptual tasks at fixation, authors report the disaggregate performance estimates for leftward, rightward, upper and lower visual fields. (Carrasco et al., 2001; Fuller and Carrasco, 2009; Zito et al., 2016a; Barbot et al., 2021; Fracasso et al., 2023). To allow a more immediate comparison between our study and other reports in the literature, we opted for including such a descriptive figure and analysis also in our manuscript.

**Figure 8.**
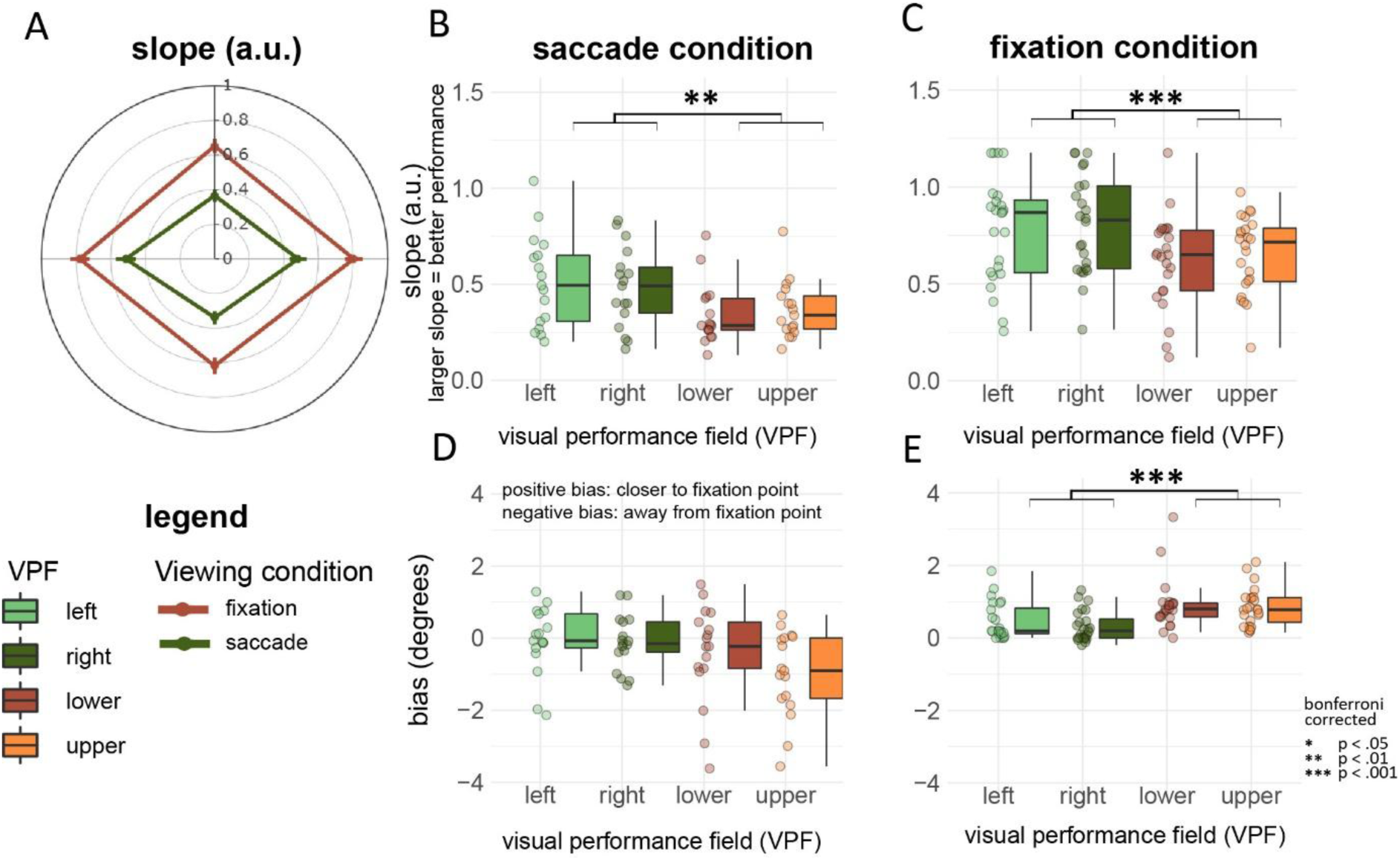
Slope and bias estimates per visual field and viewing conditions. **(A)** Polar plot displaying the slopes for all four visual fields and viewing conditions. **(B, C)** Boxplots showing the slope per visual field during the saccade and fixation condition, respectively. **(D,E)** Boxplots showing the bias per visual field during the saccade and fixation condition, respectively.

Participant performance (slope estimates) tended to be better (steeper) along the horizontal compared to the vertical meridian, with no notable differences between left and right visual fields, or upper and lower visual fields (Fig. 8a-c, statistical results reported in the figure). The same pattern of results was observed in the saccade as well as in the fixation condition, although participants were more accurate in the latter, as expected (Deubel et al., 1996).

In the saccade condition participants judged the displacement as being shifted along the direction of the first movement, which is compatible with previous findings (Matin and Pearce, 1965; Honda, 1989; Dassonville et al., 1992; Schlag and Schlag-Rey, 1995; Deubel et al., 1996; Cai et al., 1997; De Pisapia et al., 2010; Fracasso et al., 2015; Fracasso and Melcher, 2016). In contrast, the bias in the fixation condition was against the direction of the first movement, closer to the fixation point (fig. 8d, 8e). This is compatible with existing literature using different tasks, such as orientation discrimination, target detection and contrast discrimination (Musseler et al., 1999; Eggert et al., 2001; Sheth and Shimojo, 2001; Awater and Lappe, 2006; Brenner et al., 2006; Brenner et al., 2008).

Neither in the fixation nor the saccade condition participants show a difference in slope between the lower visual and upper visual field, *t*(23) = -0.73, *ns*, *p* = 0.470, and *t*(16) = -0.70, *ns*, *p* = 0.490, respectively. A Bayes Factor analysis confirmed the results obtained by the null-hypothesis testing approach, showing Bayes Factors or 0.320 and 0.360, respectively, indicating moderate to anecdotical evidence for the null hypothesis.

### Saccade reaction times and amplitudes depend on target location

In the saccade condition, we derived saccade reaction times and amplitudes for the first saccade for each of the visual fields. Saccades made towards the left visual field had an average saccade latency of 148 ms (*sd* = 54.4), with an amplitude of M = 7.75 ms, *sd* = .59. Saccades towards the right field had an average saccade latency of 152 ms (*sd* = 52.5), with an amplitude of M = 7.67 ms, *sd* = .61.

Seemingly, saccades made toward the lower visual field have a longer reaction time (M = 169, *sd* = 51.5) and slightly larger amplitude and standard deviation (M = 9.1, *sd* = 1.13) than saccades towards the upper visual field, where the average reaction time is 144 ms, (*sd* = 30.9) with an amplitude of M=8.27, *sd* = .82. This is in consensus with previous literature, where they found that saccade reaction times are faster and more accurate in the upper visual field (Schlykowa et al., 1996; Zhou and King, 2002; Hafed and Chen, 2016; Hafed and Goffart, 2020).

## Discussion

In the current study, we measured performance along different visual fields using a task which required the involvement of both the visual and oculomotor systems. Our results indicate the presence of a robust HVA effect, at fixation as well as when participants were required to perform a saccade. We did not observe signatures of VMA, neither at fixation nor during saccades.

Overall, the saccade and fixation experimental conditions showed perceptual biases in the opposite direction. During fixation, participants reported target displacement as closer to the fovea than physically presented (Fig. 6 and Fig. 8e). In the saccade condition the tendency was inverted, with participants systematically reporting target displacements as being shifted away from the fovea, indicating a bias along with saccade direction (Fig. 7 and Fig. 8d).

### Performance at fixation

At fixation, we observed better performance along the horizontal compared to the vertical meridian (HVA), but we did not observe an asymmetry along the vertical meridian (VMA), as no difference in performance was observed between the upper and lower visual fields. Seemingly, the anisotropies generally observed at fixation on perceptual discrimination tasks (‘what am I looking at’) only partially transfer to target displacement tasks (‘where did the target move’). (Carrasco et al., 2001; Fuller and Carrasco, 2009; Zito et al., 2016; Barbot et al., 2021).

As far a response bias is concerned, at fixation participants systematically reported displacements towards the centre of the screen (an inward bias). The displacement was perceived as being closer to the starting fixation point than it physically was. This is compatible with existing literature measuring perceived position of peripheral targets (Musseler et al., 1999; Eggert et al., 2001; Sheth and Shimojo, 2001; Awater and Lappe, 2006; Brenner et al., 2006; Brenner et al., 2008).

Sheth and Shimojo (2001) used a pointing task to assess the perceived position in the periphery (∼25 dva). Participants had to retain in memory (∼2 seconds) the position of a briefly (30 ms) flashed target. The authors observed an inward bias interpreted as a visual memory effect, induced by a distorted spatial working memory in the presence of visual references (Sheth and Shimojo, 2001).

A similar account for this inward bias of perceived position has been proposed by Eggert et al. (2001) where the inward bias has been linked to a mismatch between egocentric location - retinal position of the target - and exocentric location -veridical position of the target -mapping the displaced target on the memorized internal representation of the target position (Eggert et al., 2001).

In these examples, authors measured perceived position, and not displacement, as in the current study. Moreover, the experimental designs incorporated a significant delay between target presentation and the participant response phase. In these conditions an account based on a corrupted memory trace could be valid for measured inward biases of perceived position, however the same cannot be applied to our results. In our case, there was no delay or gap between the target and the displaced stimuli. Moreover, there was a clear motion cue (displacement), indicating the direction of the displacement. If participants relied on motion cues alone, no bias should have been observed. On the contrary, we observed a systematic inward bias at fixation; this result suggests that such bias might be mediated by low-level localization mechanisms that occur prior to motion integration in the visual periphery (Burr and Santoro, 2001). Interestingly, a similar inward bias can be observed for stimuli presented in the periphery as close as ∼3 dva (Brenner et al., 2008), and positively scales with eccentricity (Brenner et al., 2006).

We observed that the inward bias during fixation is stronger along the vertical meridian compared to the horizontal meridian. For targets located in the upper or lower visual field, the displacement was perceived closer to the fixation point than for targets located across the horizontal meridian. We speculate that this asymmetry between the horizontal and vertical inward bias could be linked to polar angle asymmetries observed in early visual cortex (Himmelberg et al., 2023).

### Individual differences in response bias during saccades

We performed a clustering analysis to exclude participants whose responses were characterized by a strong response bias. These participants had a strong bias in reporting the direction of the displacement along the direction of first movement, regardless of the physical direction of the displacement (Deubel et al., 1996). The strength of the bias in these participants was such that it prevented any reliable attempt at estimating its magnitude, other than via extrapolation. On the other hand, the same participants could successfully discriminate the direction of the displacement at fixation, thus we speculate that the nature of the outward bias observed in the saccade condition was not perceptual in nature, but rather linked to the response: it is likely that these participants gave their response following a heuristic based solely on the direction of the first movement, that is: saccade direction. It is possible that the participant might have misunderstood the given instruction, or it could be argued that we used a too narrow stimulus range, however we believe this is unlikely, as the biased judgements in the saccade condition did not transfer to the fixation condition. Given the pattern of results, we believe it is more likely that these participants either could not discriminate displacement direction or had little confidence in their discrimination ability and hence relied on a simply reporting the direction of the eye movement.

We removed these participants due to the unreliable estimates in the saccade condition. After removing these participants, a general bias along the first movement was still evident, as reported in the literature (Deubel et al., 1996), although in this case we could reliably estimate its magnitude without relying on extrapolation.

### Performance during saccades

We observed the known HVA in the saccade condition, with better performance along the horizontal meridian compared to the vertical meridian, which is consistent with previous findings reported at fixation (Carrasco et al., 2001; Kristjansson and Sigurdardottir, 2008; Barbot et al., 2021).

Again, we did not observe a difference in performance between the upper and lower visual field during the saccade condition, although a trend compatible with the existing literature at fixation was present. It could be argued that our design lacked the sensitivity to detect differences in precision (slope) between the upper and lower visual field condition. However, we consider this option unlikely, as we successfully detected differences between the horizontal and vertical conditions. These findings indicate that our participants’ performance was neither at ceiling, nor at floor. Additionally, individual slope estimates varied substantially across participants (Figure. 7c).

In contrast with the inward bias found at fixation, during the saccade condition participants systematically judged the displacement along the direction of the first movement. We also observed a trend in bias estimates between visual fields: the bias in the vertical condition tended to be larger compared to the horizontal condition, and bias in upper visual field tended to be larger than in lower VF condition. However, it is important to note that these trends did not survive Bonferroni corrections. If we were to speculate on these trends, it is interesting to highlight that the pattern is compatible with the HVA and VMA observed at fixation in perceptual discrimination (Carrasco et al., 2001; Fuller and Carrasco, 2009; Zito et al., 2016; Barbot et al., 2021), and goes in the opposite direction compared to the perisaccadic modulation shown in Fracasso et al. (2023).

The outward bias during the saccade condition is in consensus with previous findings in mislocalization literature and SSID (Matin and Pearce, 1965; Honda, 1989; Dassonville et al., 1992; Schlag and Schlag-Rey, 1995; Deubel et al., 1996; Cai et al., 1997; De Pisapia et al., 2010; Fracasso et al., 2015; Fracasso and Melcher, 2016). In mislocalization studies, participants are required to report the perceived location of briefly presented flashes around saccade onset, whereas in SSID participants are required to report the perceived displacement of the saccade target.

Mislocalization studies generally report shifts along the direction of the saccade for flashes presented between the starting and landing fixation points (Matin and Pearce, 1965; Honda, 1989; Dassonville et al., 1992; Schlag and Schlag-Rey, 1995; Deubel et al., 1996; Cai et al., 1997), and compression against saccade direction for targets presented beyond the landing fixation point (Ross et al., 1997; VanRullen, 2004; Awater and Lappe, 2006; Shim and Cavanagh, 2006; Ostendorf et al., 2007; Buonocore and Melcher, 2015). It has been shown that compression is strongly linked to the presence of visual references (Krekelberg et al., 2000; Lappe et al., 2000).

On the other hand, SSID studies (including the current study) report a general outward bias, along the direction of the eye movement (Dassonville et al., 1992; Deubel et al., 1996; Deubel et al., 1998). It is important to note that models of perceived mislocalization successfully predict patterns of shifts and compression (VanRullen, 2004), but predict an absence of mislocalization corresponding to the saccade target itself, that is: briefly flashed probes around the saccadic target should be perceived veridically. SSID results indicate a different pattern, showing that the saccade target itself can be strongly misperceived as being shifted along the saccade direction. Another study supports this finding: in preparation of a saccade, Zirnsak and Colleagues (2014) showed that receptive fields shift towards the saccadic target, leading to mislocalized presaccadic stimuli as being nearer to the saccadic target, along with the direction of the saccade (Zirnsak et al., 2014).

### Horizontal-meridian-asymmetry and vertical-meridian-asymmetry in the saccade condition

We observed the classic horizontal-vertical asymmetry for saccade and fixation in SSID, while we did not observe robust signs of the vertical meridian asymmetry.

The vertical meridian asymmetry is generally observed at fixation in orientation discrimination and related perceptual tasks (Carrasco et al., 2001; Kristjansson and Sigurdardottir, 2008; Barbot et al., 2021), moreover it has also been reported in the context of deployment of attentional resources (He et al., 1996), and visual illusion (Rubin et al., 1996). The task adopted in this investigation (SSID) is different compared to the tasks in which the VMA is normally observed. In SSID the focus is on the displacement of target in space (‘where did the target move?’), instead of perceptual discrimination per se (‘what I am looking at’).

It could be speculated that these tasks rely on distinct visual processes or pathways, probing different streams of visual information: the ventral ‘what’ stream and dorsal ‘where’ stream (Goodale and Milner, 1992). Differences in visual perception along the vertical meridian might serve a functional role in answering the question ‘what am I looking at?’, in which it is often more important to identify what appears in our lower visual field than in the upper visual field (e.g. grasping for objects, avoiding falls, avoiding falls, or stepping over hazards like a snake). In contrast, vertical meridian asymmetries (VMA) may not extend to tasks involving spatial displacement, such as determining where an object moved (Previc, 1990).

Alternatively, these results could arise from the interaction between known neural anisotropies between the superior colliculus and primary visual cortex. In primary visual cortex a larger portion of surface area is dedicated to processing the horizontal versus the vertical meridian and the lower compared to the upper visual field (Himmelberg et al., 2022; Kupers et al., 2022; Himmelberg et al., 2023), whereas the opposite has been observed in the SC (Hafed and Chen, 2016). One might speculate about a compensatory mechanism balancing between opposite anisotropies in tasks relying heavily on both the perceptual and the oculomotor system, such as SSID. Visual perception during saccades could be dominated by the SC, an ontogenetically older structure, present since the early development of mammals (White et al., 2017; Beltramo and Scanziani, 2019). The SC is more sensitive for stimuli presented in the upper compared to the lower visual field, possibly to serve early detection of dangers from above, for example flying predators. Literature shows that UV contrast is predominantly processed in the upper visual field in mice, which helps detect predators in the sky (Qiu et al., 2021). This interpretation is corroborated by recent evidence from monkey neurophysiology (Hafed and Chen, 2016). Moreover, it has recently been shown how orientation discrimination accuracy can be comparable between the upper and lower visual fields in close proximity with eye movements (Liu et al., 2024), or even show an advantage towards the upper visual field in the perisaccadic interval (Fracasso et al., 2023). This hypothesis assumes that the modes of operation governing the perception-action cycle differs between fixation and during (or in the vicinity of) saccades. It has been proposed that visual perception at fixation evolved to support the motor system when performing precise actions towards the lower visual field, within the peri-personal space (Previc, 1990), for example: catching small prey, or grasping and reaching for objects. These actions heavily rely on visual feedback (Khan and Lawrence, 2005; Kristjansson and Sigurdardottir, 2008; Rossit et al., 2013; Gottwald et al., 2015).

As a third alternative, the upper visual field advantage observed behaviourally could be linked to biases introduced while scanning the environment during natural locomotion, requiring frequent and rapid orienting movements predominantly towards the upper visual field (Matthis et al., 2018; Muller et al., 2023). Please note that these hypotheses are not mutually exclusive.

### Concluding remarks

We measured performance along different visual fields with SSID, a task which required the involvement of both the visual and oculomotor systems. Our results indicate the presence of a robust HVA effect, at fixation as well as when participants were required to perform a saccade, in the absence of a robust VMA (although we observed a trend in bias estimates).

This pattern of results is consistent with current literature on behavioural anisotropies measured in the context of saccades, showing robust HMA and an overall reduction in VMA. We proposed different non-mutually exclusive hypotheses to account for this pattern of results, as different mechanisms can interact and contribute to the observed effects.

## Statements and Declarations

Nothing to disclose.

## Acknowledgements

A.F. was supported by a grant from the Biotechnology and Biology Research Council (BBSRC, grant number: BB/S006605/1) and the Bial Foundation (Bial Foundation Grants Program; Grant id: A-29315, number: 203/2020, grant edition: G-15516). A.B. was supported by a grant from the Italian Minister of Research and University (PRIN 2022_PNRR-P2022ST78T).

## Data availability

original data will be made available on reasonable request.

## References

Awater H, Lappe M (2006) Mislocalization of perceived saccade target position induced by perisaccadic visual stimulation. The Journal of neuroscience : the official journal of the Society for Neuroscience 26:12–20.

Barbot A, Xue S, Carrasco M (2021) Asymmetries in visual acuity around the visual field. Journal of vision 21:2.

Beltramo R, Scanziani M (2019) A collicular visual cortex: Neocortical space for an ancient midbrain visual structure. Science (New York, NY) 363:64–69.

Brainard DH, Vision S (1997) The psychophysics toolbox. Spatial vision 10:433–436.

Brenner E, Mamassian P, Smeets JB (2008) If I saw it, it probably wasn’t far from where I was looking. Journal of vision 8:7–7.

Brenner E, van Beers RJ, Rotman G, Smeets JB (2006) The role of uncertainty in the systematic spatial mislocalization of moving objects. Journal of Experimental Psychology: Human Perception and Performance 32:811.

Bridgeman B, Hendry D, Stark L (1975) Failure to detect displacement of the visual world during saccadic eye movements. Vision research 15:719–722.

Brunner G, Gajwani R, Gross J, Gumley A, Timmerman RH, Taylor R, Krishnadas R, Lawrie SM, Schwannauer M, Schultze-Lutter F (2025) Choroid plexus morphology in schizophrenia and early-stage psychosis: A cross-sectional study. Schizophrenia research 275:107–114.

Buonocore A, Melcher D (2015) Interference during eye movement preparation shifts the timing of perisaccadic compression. Journal of vision 15:3.

Buonocore A, Fracasso A, Melcher D (2017) Pre-saccadic perception: Separate time courses for enhancement and spatial pooling at the saccade target. PloS one 12:e0178902.

Burr DC, Santoro L (2001) Temporal integration of optic flow, measured by contrast and coherence thresholds. Vision research 41:1891–1899.

Cai RH, Pouget A, Schlag-Rey M, Schlag J (1997) Perceived geometrical relationships affected by eye- movement signals. Nature 386:601–604.

Carrasco M, Talgar CP, Cameron EL (2001) Characterizing visual performance fields: effects of transient covert attention, spatial frequency, eccentricity, task and set size. Spatial vision 15:61–75.

Cornelissen FW, Peters EM, Palmer J (2002) The Eyelink Toolbox: eye tracking with MATLAB and the Psychophysics Toolbox. Behavior Research Methods, Instruments, & Computers 34:613–617.

Dalby C, Dibble A, Carvalheiro J, Queirazza F, Sevegnani M, Harvey M, Svanera M, Fracasso A (2024) Delineating In-Vivo T1-Weighted Intensity Profiles Within the Human Insula Cortex Using 7- Tesla MRI. bioRxiv:2024.2008. 2005.605123.

Dassonville P, Schlag J, Schlag-Rey M (1992) Oculomotor localization relies on a damped representation of saccadic eye displacement in human and nonhuman primates. Visual neuroscience 9:261–269.

De Pisapia N, Kaunitz L, Melcher D (2010) Backward masking and unmasking across saccadic eye movements. Current biology : CB 20:613–617.

Deubel H, Schneider WX, Bridgeman B (1996) Postsaccadic target blanking prevents saccadic suppression of image displacement. Vision research 36:985–996.

Deubel H, Bridgeman B, Schneider WX (1998) Immediate post-saccadic information mediates space constancy. Vision research 38:3147–3159.

Deubel H, Schneider WX, Bridgeman B (2002) Transsaccadic memory of position and form. Progress in brain research 140:165–180.

Eggert T, Ditterich J, Straube A (2001) Mislocalization of peripheral targets during fixation. Vision research 41:343–352.

Fabius JH, Fracasso A, Van der Stigchel S (2016) Spatiotopic updating facilitates perception immediately after saccades. Scientific reports 6:34488.

Fabius JH, Fracasso A, Nijboer TCW, Van der Stigchel S (2019) Time course of spatiotopic updating across saccades. Proceedings of the National Academy of Sciences of the United States of America 116:2027–2032.

Fabius JH, Fracasso A, Acunzo DJ, Van der Stigchel S, Melcher D (2020) Low-Level Visual Information Is Maintained across Saccades, Allowing for a Postsaccadic Handoff between Visual Areas. The Journal of neuroscience : the official journal of the Society for Neuroscience 40:9476–9486.

Fracasso A, Melcher D (2016) Saccades Influence the Visibility of Targets in Rapid Stimulus Sequences: The Roles of Mislocalization, Retinal Distance and Remapping. Frontiers in systems neuroscience 10:58.

Fracasso A, Caramazza A, Melcher D (2010) Continuous perception of motion and shape across saccadic eye movements. Journal of vision 10:14.

Fracasso A, Kaunitz L, Melcher D (2015) Saccade kinematics modulate perisaccadic perception. Journal of vision 15.

Fracasso A, Buonocore A, Hafed ZM (2023) Peri-saccadic orientation identification performance and visual neural sensitivity are higher in the upper visual field. Journal of Neuroscience 43:6884–6897.

Fracasso A, Targher S, Zampini M, Melcher D (2013) Fooling the eyes: the influence of a sound-induced visual motion illusion on eye movements. PloS one 8:e62131.

Fuller S, Carrasco M (2009) Perceptual consequences of visual performance fields: the case of the line motion illusion. Journal of vision 9:13.11-17.

Gelman A, Hill J (2007) Data analysis using regression and multilevel/hierarchical models: Cambridge university press.

Goodale MA, Milner AD (1992) Separate visual pathways for perception and action. Trends in neurosciences 15:20–25.

Gottwald VM, Lawrence GP, Hayes AE, Khan MA (2015) Representational momentum reveals visual anticipation differences in the upper and lower visual fields. Experimental brain research 233:2249–2256.

Grujic N, Brehm N, Gloge C, Zhuo W, Hafed ZM (2018) Perisaccadic perceptual mislocalization is different for upward saccades. Journal of neurophysiology 120:3198–3216.

Hafed ZM (2018) Superior Colliculus: A Vision for Orienting. Current biology : CB 28:R1111–r1113.

Hafed ZM, Chen CY (2016) Sharper, Stronger, Faster Upper Visual Field Representation in Primate Superior Colliculus. Current biology : CB 26:1647–1658.

Hafed ZM, Goffart L (2020) Gaze direction as equilibrium: more evidence from spatial and temporal aspects of small-saccade triggering in the rhesus macaque monkey. Journal of neurophysiology 123:308–322.

Hafed ZM, Hoffmann KP, Chen CY, Bogadhi AR (2023) Visual Functions of the Primate Superior Colliculus. Annual review of vision science.

He S, Cavanagh P, Intriligator J (1996) Attentional resolution and the locus of visual awareness. Nature 383:334–337.

Himmelberg MM, Winawer J, Carrasco M (2022) Linking individual differences in human primary visual cortex to contrast sensitivity around the visual field. Nature communications 13:3309.

Himmelberg MM, Winawer J, Carrasco M (2023) Polar angle asymmetries in visual perception and neural architecture. Trends in neurosciences 46:445–458.

Honda H (1989) Perceptual localization of visual stimuli flashed during saccades. Perception & psychophysics 45:162–174.

Kaunitz L, Fracasso A, Melcher D (2011) Unseen complex motion is modulated by attention and generates a visible aftereffect. Journal of vision 11:10.

Kaunitz L, Fracasso A, Lingnau A, Melcher D (2013) Non-conscious processing of motion coherence can boost conscious access. PloS one 8:e60787.

Kaunitz LN, Fracasso A, Skujevskis M, Melcher D (2014) Waves of visibility: probing the depth of inter-ocular suppression with transient and sustained targets. Frontiers in psychology 5:804.

Khan MA, Lawrence GP (2005) Differences in visuomotor control between the upper and lower visual fields. Experimental brain research 164:395–398.

Krekelberg B, Lappe M, Whitney D, Cavanagh P, Eagleman DM, Sejnowski TJ (2000) The position of moving objects. Science (New York, NY) 289:1107a.

Kristjansson A, Sigurdardottir HM (2008) On the benefits of transient attention across the visual field. Perception 37:747–764.

Kupers ER, Benson NC, Carrasco M, Winawer J (2022) Asymmetries around the visual field: From retina to cortex to behavior. PLoS computational biology 18:e1009771.

Lappe M, Awater H, Krekelberg B (2000) Postsaccadic visual references generate presaccadic compression of space. Nature 403:892–895.

Lee MD, Wagenmakers E-J (2014) Bayesian cognitive modeling: A practical course: Cambridge university press.

Liu X, Melcher D, Carrasco M, Hanning NM (2024) Presaccadic preview shapes postsaccadic processing more where perception is poor. Proceedings of the National Academy of Sciences of the United States of America 121:e2411293121.

Matin L, Pearce DG (1965) Visual Perception of Direction for Stimuli Flashed During Voluntary Saccadic Eye Movements. Science (New York, NY) 148:1485–1488.

Matthis JS, Yates JL, Hayhoe MM (2018) Gaze and the control of foot placement when walking in natural terrain. Current Biology 28:1224–1233. e1225.

Melcher D, Fracasso A (2012) Remapping of the line motion illusion across eye movements. Experimental brain research 218:503–514.

Mohler CW, Wurtz RH (1976) Organization of monkey superior colliculus: intermediate layer cells discharging before eye movements. Journal of neurophysiology 39:722–744.

Morey RD, Rouder JN, Jamil T, Morey MRD (2015) Package ‘bayesfactor’. In.

Muller KS, Matthis J, Bonnen K, Cormack LK, Huk AC, Hayhoe M (2023) Retinal motion statistics during natural locomotion. eLife 12:e82410.

Musseler J, van der Heijden AH, Mahmud SH, Deubel H, Ertsey S (1999) Relative mislocalization of briefly presented stimuli in the retinal periphery. Perception & psychophysics 61:1646–1661.

Ostendorf F, Fischer C, Finke C, Ploner CJ (2007) Perisaccadic compression correlates with saccadic peak velocity: differential association of eye movement dynamics with perceptual mislocalization patterns. The Journal of neuroscience : the official journal of the Society for Neuroscience 27:7559–7563.

Previc FH (1990) Functional specialization in the lower and upper visual fields in humans: Its ecological origins and neurophysiological implications. Behavioral and Brain Sciences 13:519–542.

Qiu Y, Zhao Z, Klindt D, Kautzky M, Szatko KP, Schaeffel F, Rifai K, Franke K, Busse L, Euler T (2021) Natural environment statistics in the upper and lower visual field are reflected in mouse retinal specializations. Current biology : CB 31:3233–3247 e3236.

Ross J, Morrone MC, Burr DC (1997) Compression of visual space before saccades. Nature 386:598–601.

Rossit S, McAdam T, McLean DA, Goodale MA, Culham JC (2013) fMRI reveals a lower visual field preference for hand actions in human superior parieto-occipital cortex (SPOC) and precuneus. Cortex; a journal devoted to the study of the nervous system and behavior 49:2525–2541.

Rubin N, Nakayama K, Shapley R (1996) Enhanced perception of illusory contours in the lower versus upper visual hemifields. Science (New York, NY) 271:651–653.

Schlag J, Schlag-Rey M (1995) Illusory localization of stimuli flashed in the dark before saccades. Vision research 35:2347–2357.

Schlykowa L, Hoffmann KP, Bremmer F, Thiele A, Ehrenstein WH (1996) Monkey saccadic latency and pursuit velocity show a preference for upward directions of target motion. Neuroreport 7:409–412.

Schweitzer R, Rolfs M (2020) An adaptive algorithm for fast and reliable online saccade detection. Behavior research methods 52:1122–1139.

Sheth BR, Shimojo S (2001) Compression of space in visual memory. Vision research 41:329–341.

Shim WM, Cavanagh P (2006) Bi-directional illusory position shifts toward the end point of apparent motion. Vision research 46:3214–3222.

Taylor R, Buonocore A, Fracasso A (2024) Saccadic “inhibition” unveils the late influence of image content on oculomotor programming. Experimental brain research 242:2281–2294.

VanRullen R (2004) A simple translation in cortical log-coordinates may account for the pattern of saccadic localization errors. Biological cybernetics 91:131–137.

White BJ, Kan JY, Levy R, Itti L, Munoz DP (2017) Superior colliculus encodes visual saliency before the primary visual cortex. Proceedings of the National Academy of Sciences 114:9451–9456.

Wurtz RH, Goldberg ME (1972) The role of the superior colliculus in visually-evoked eye movements. Bibliotheca ophthalmologica : supplementa ad ophthalmologica 82:149–158.

Wurtz RH, Albano JE (1980) Visual-motor function of the primate superior colliculus. Annual review of neuroscience 3:189–226.

Zhou W, King WM (2002) Attentional sensitivity and asymmetries of vertical saccade generation in monkey. Vision research 42:771–779.

Zirnsak M, Steinmetz NA, Noudoost B, Xu KZ, Moore T (2014) Visual space is compressed in prefrontal cortex before eye movements. Nature 507:504–507.

Zito GA, Cazzoli D, Muri RM, Mosimann UP, Nef T (2016) Behavioral Differences in the Upper and Lower Visual Hemifields in Shape and Motion Perception. Frontiers in behavioral neuroscience 10:128.

